# Loss of *Cnot6l* impairs inosine RNA modifications in mouse oocytes

**DOI:** 10.1101/2020.11.04.358010

**Authors:** Pavla Brachova, Nehemiah S. Alvarez, Lane K. Christenson

**Affiliations:** Department of Molecular and Integrative Physiology, University of Kansas Medical Center, Kansas City, KS 66160

## Abstract

Mammalian oocytes must degrade maternal transcripts through a process called translational mRNA decay, in which maternal mRNA undergoes translational activation, followed by deadenylation and mRNA decay. Once a transcript is translationally activated, it becomes deadenylated by the CCR4-NOT complex. Knockout of *Cnot6l*, a deadenylase within the CCR4-NOT complex, results in mRNA decay defects during MI entry. Knockout of *Btg4*, an adaptor protein of the CCR4-NOT complex, results in mRNA decay defects following fertilization. Therefore, mechanisms controlling mRNA turnover have significant impacts on oocyte competence and early embryonic development. Post-transcriptional inosine RNA modifications can impact mRNA stability, possibly through a translation mechanism. Here, we assessed inosine RNA modifications in oocytes from *Cnot6l^-/-^* and *Btg4^-/-^* mice, which display stabilization of mRNA and over-translation of the stabilized transcripts. If inosine modifications have a role in modulating RNA stability, we hypothesize that in these mutant backgrounds, we would observe changes or a disruption in inosine mRNA modifications. To test this, we used a computational approach to identify inosine RNA modifications in total and polysomal RNA-seq data during meiotic maturation (GV, MI, and MII stages). We observed pronounced depletion of inosine mRNA modifications in oocytes from *Cnot6l^-/-^*, but not in *Btg4^-/-^* mice. Additionally, analysis of ribosome-associated RNA revealed clearance of inosine modified mRNA. These observations suggest a novel mechanism of mRNA clearance during oocyte maturation, in which inosine-containing transcripts decay in an independent, but parallel mechanism to CCR4-NOT deadenylation.

## Introduction

Growing mammalian oocytes transcribe and store transcripts that support meiotic maturation, fertilization, and early embryo development (Bachvarova and De Leon 1980; Bachvarova 1985; De La Fuente et al. 2004; De La Fuente and Eppig 2001). The transition of maternal to zygotic control of development is termed the maternal to zygotic transition (MZT), and occurs in the absence of transcription (Piko and Clegg 1982; De La Fuente and Eppig 2001). As oocytes resume meiosis, a large-scale, but selective wave of maternal transcripts is targeted for translation and subsequent degradation (Su et al. 2007; Svoboda et al. 2015; Sha et al. 2019) in a process called translational mRNA decay. By the 2-cell stage of mouse early embryogenesis, approximately 90% of maternal RNA are degraded (Schellander et al. 2007). The activation of translation requires specific 3’ UTR elements, such as polyadenylation signals and cytoplasmic polyadenylation elements (Chen et al. 2011; Dai et al. 2019). Once transcripts undergo translational activation following the lengthening of their poly(A) tails, they are subject to decay by deadenylation (Ozturk and Uysal 2017). CCR4-NOT is the major, multi-subunit deadenylase complex involved in poly(A) shortening (Collart and Panasenko 2012). A functional CCR4-NOT complex is essential for the proper decay of maternal oocyte mRNA, and for the development of a fertilized oocyte into an embryo (Sha et al. 2018; Horvat et al. 2018; Yu et al. 2016; Dumdie et al. 2018).

An understanding of the individual components of the CCR4-NOT complex has illuminated specific features of mRNA transcripts leading to their specific and selective decay. The ribonuclease *Cnot6l* is uniquely expressed in mouse oocytes, and regulates deadenylation of transcripts undergoing translational mRNA decay during oocyte maturation (Sha et al. 2018; Horvat et al. 2018; Vieux and Clarke 2018). *Cnot6l^-/-^* mice are severely subfertile due to the overtranslation of undegraded maternal transcripts, resulting in spindle defects during the completion of meiosis (Sha et al. 2018; Horvat et al. 2018). The CCR4-NOT deadenylase complex also relies on the adaptor RNA binding proteins, ZFP36L2 and BTG4, to associate with cytoplasmic transcripts (Ball et al. 2014; Dumdie et al. 2018; Doidge et al. 2012; Liu et al. 2016; Wu and Dean 2016; Pasternak et al. 2016). *Zfp36l2^-/-^* mice are infertile due to decreased oocyte numbers as well as decreased rates of oocyte maturation (Ball et al. 2014; Dumdie et al. 2018). *Btg4^-/-^* mice are also infertile due to embryonic arrest at the 2-cell and 4-cell stages (Yu et al. 2016; Sha et al. 2018). Cumulatively, these models demonstrate that the translational mRNA decay machinery is necessary to establish the correct dosage of RNA during the MZT, and dysfunction or loss of critical components within the mRNA degradation pathway leads to infertility.

Previous work has established inosine post-transcriptional RNA modifications can impact mRNA stability through a translation mechanism (Licht et al. 2019). Since the *Cnot6l^-/-^* and *Btg4^-/-^* mice result in stabilization of mRNA and overtranslation of stabilized transcripts, we hypothesized that in these mutant backgrounds, we would observe an increase in inosine modifications in mRNA. Surprisingly, we observed a significant decrease in inosine modifications in oocytes from *Cnot6l^-/-^*, but not in *Btg4^-/-^* mice, suggesting that in the absence of *Cnot6l*, mRNA degradation of inosine-modified transcripts is still occurring, while unmodified transcripts are stabilized. Our data suggest novel components of translational mRNA decay during oocyte maturation.

## Results

### Inosine RNA modifications are lost in oocytes from Cnot6l^-/-^ mice

Using our computational approach (Brachova et al. 2019), we identified transcriptome wide inosine RNA modifications (Supplemental Fig. S1) in previously reported RNA-seq datasets (Sha et al. 2018) from wild-type (WT), *Cnot6l^-/-^, Btg4^-/-^* oocytes at 0, 8, and 16 h after in vitro maturation. These three time points correspond to the GV, MI, and MII oocyte stages, respectively. The number of unique transcripts with inosine RNA modifications in *Cnot6l^-/-^* oocytes (134 ± 28 in GV, 159 ± 0 in MI, 163 ± 3 in MII, and 160 ± 7 in Zyg; Mean ± SEM) were significantly lower than both WT oocytes and *Btg4^-/-^* oocytes at all oocyte/embryo stages (WT: 1750 ± 337 in GV, 999 ± 109 in MI, 1132 ± 213 in MII, and 1311 ± 562 in Zyg; *Btg4^-/-^*: 1749 ± 197 in GV, 1412 ± 30 in MI, 1579 ± 103 in MII, and 1441 ± 200 in Zyg; one-way ANOVA; Fig. 1a). No difference in number of transcripts with inosine RNA modifications was detected between WT and *Btg4^-/-^* oocytes at any stage (Fig. 1a). To confirm that the reduction in transcripts with inosine RNA modifications in *Cnot6l^-/-^oocytes* was not due to a differential pattern of RNA expression, we identified inosine modifications only in transcripts found in all samples across all groups (8,686 transcripts, total RNA; 6,006 transcripts, ribosome-associated transcripts). The results (Fig. 1c), indicate that the loss of inosine RNA modification was not due to differential gene expression.

**Figure 1.**
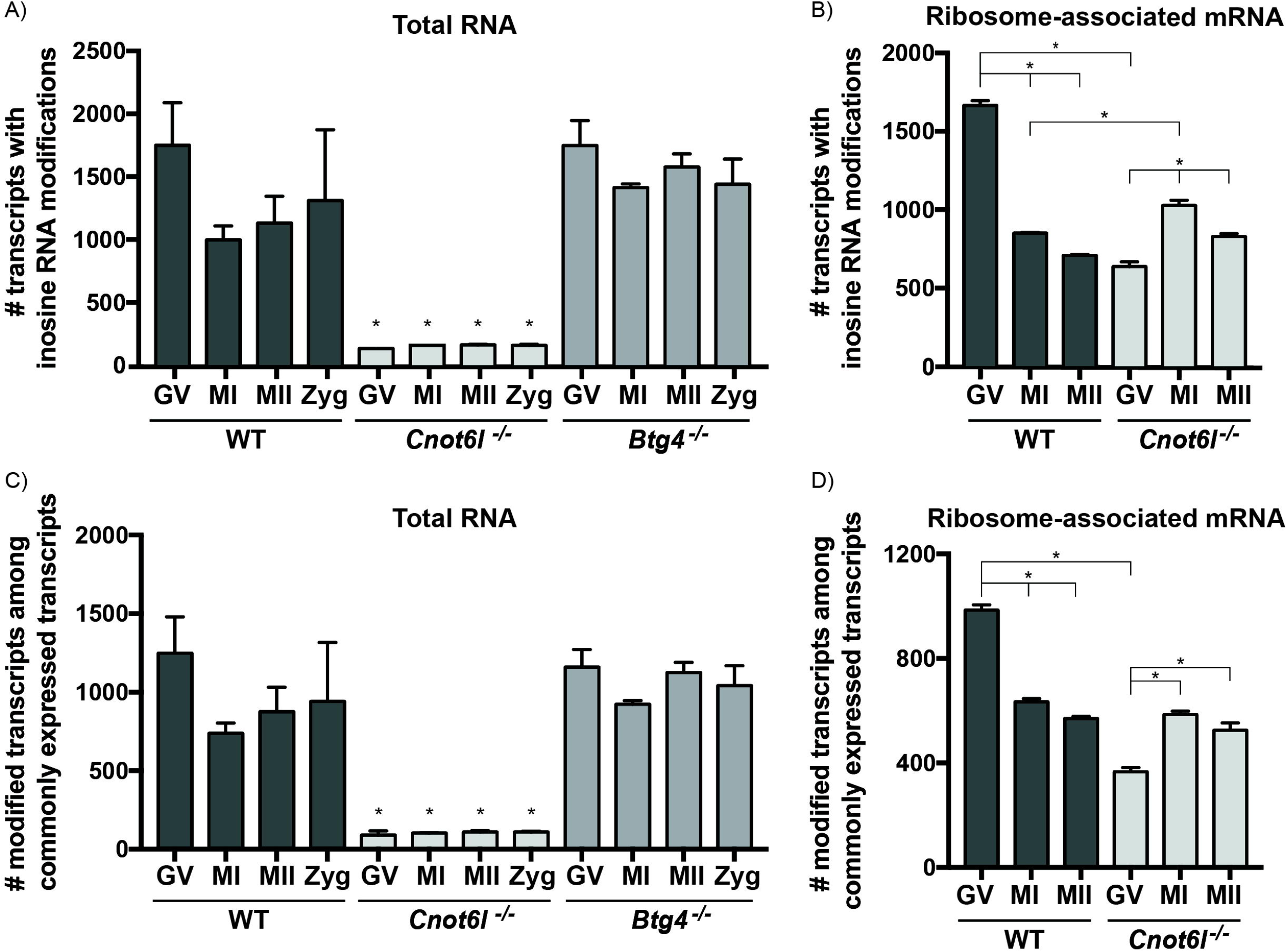
Inosine RNA modifications occur in transcripts enriched in the ribosome-associated RNA. A-B) The number of unique inosine modified transcripts identified in WT, *Cnot6l^-/-^*, and *Btg4^-/-^* GV, MI, MII, and Zyg in total RNA (A) and ribosome-associated mRNA (B). C-D) Among common transcripts present in all samples, the number of unique inosine modified transcripts identified in WT, *Cnot6l^-/-^*, and *Btg4^-/-^* GV, MI, MII, and Zyg in total RNA (C) and ribosome-associated mRNA (D).

### Inosine RNA modifications are enriched in ribosome-associated mRNA

Our previous study linked inosine RNA modifications to the process of translation through RNA modifications of codons (Brachova et al. 2019). To test if the accumulated and over-translated maternal transcripts found in oocytes of *Cnot6l^-/-^* mice were enriched in inosine RNA modifications, we cataloged inosine RNA modifications in ribosome-associated mRNA at the GV, MI, and MII stages in WT and *Cnot6l^-/-^* oocytes. In the ribosome-associated mRNA of WT GV oocytes, the number of inosine-modified transcripts decreased during oocyte maturation, possibly indicating clearance of inosine-modified transcripts (Fig. 1b). Conversely, in *Cnot6l^-/-^* oocytes, significantly fewer transcripts are inosine-modified at the GV stage, and the number of inosine-modified transcripts increases, rather than decreases during oocyte maturation (Fig. 1b). Even among commonly expressed transcripts, the pattern is similar (Fig. 1d). Our data show that globally, the total RNA fraction of *Cnot6l^-/-^* oocytes have fewer inosine modified transcripts, however, these modified transcripts are still detected in the ribosome-associated fraction.

### Pattern of inosine RNA modifications in total and ribosome-associated mRNA

In addition to having the fewest inosine modified transcripts, oocytes from *Cnot6l^-/-^* mice had the lowest proportion of the transcriptome that was modified (GV: 1.0 ± 0.3%, MI: 1.2 ± 0.02%, MII:1.2 ± 0.03%, Zyg: 1.3 ± 0.05%), compared to WT oocytes (GV: 14.4 ± 2.6%, MI: 8.5 ± 0.7%, MII: 10.1 ± 1.7%, Zyg: 10.9 ± 4.3%) and *Btg4^-/-^* oocytes (GV: 13.4 ± 1.2%, MI: 10.6 ± 0.3%, MII: 13.0 ± 0.7%, Zyg: 12.0 ± 1.4%; X^2^ p<0.5; Fig. 2a). Among ribosome-associated mRNA, a lower proportion of transcripts were modified in *Cnot6l^-/-^* oocytes (GV: 6.1 ± 0.2%, MI: 9.7 ± 0.2%, MII: 8.7 ± 0.4%,), compared to WT oocytes (GV: 16.4 ± 0.3%, MI: 10.6 ± 0.2%, MII: 9.5 ± 0.1%, Fig. 2b). Comparing the number of identified inosine RNA modifications per transcript, on average, WT oocytes and *Btg4*^-/-^ oocytes had similar patterns, while *Cnot6l*^-/-^ oocytes contained transcripts with fewer modifications per transcript (blue and green lines, Fig. 2c). However, within ribosome-associated mRNA, the pattern of inosine RNA modifications per transcript was similar (Fig. 2c). To further characterize the nature of inosine RNA modifications in oocytes from WT, *Cnot6l^-/-^*, and *Btg4^-/-^* mice, the location of modifications within protein-coding genes was determined by annotating the location of each inosine RNA modification (5’ UTR, coding, intron, and 3’ UTR). Overall, the distribution of inosine RNA modifications is similar between all stages of oocyte maturation in WT and *Btg4^-/-^* mice, with the majority of modifications occurring in the CDS and 3’ UTR. Conversely, oocytes from *Cnot6l^-/-^* mice display a higher proportion of intron inosine RNA modifications (Fig. 2d). Within transcripts from ribosome-associated mRNA, the pattern of inosine RNA modifications was similar (Fig. 2d).

**Figure 2.**
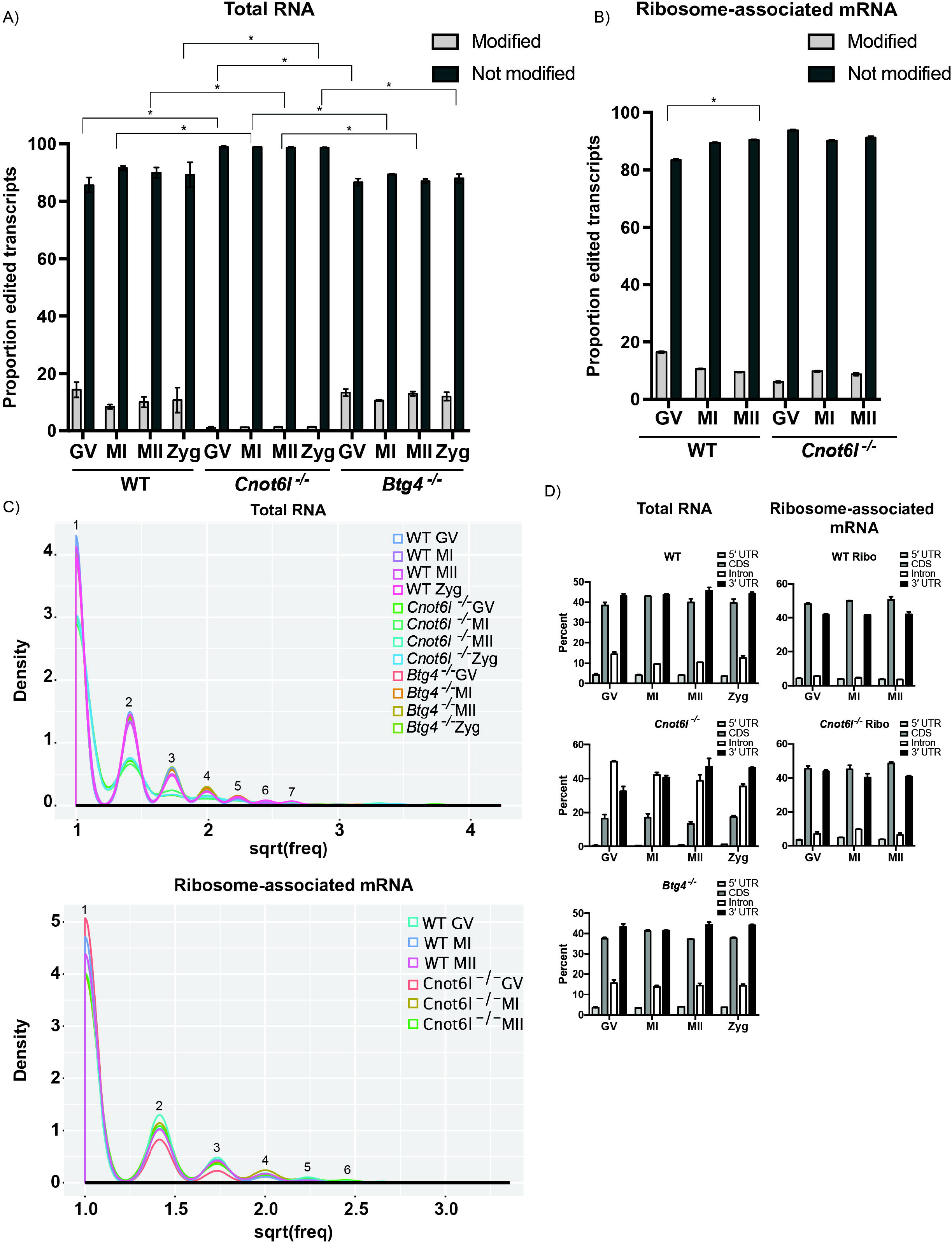
Distinct pattern of inosine RNA modifications from WT, *Cnot6l^-/-^*, and *Btg4^-/-^* oocytes and zygotes. A-B) Proportion of the transcriptome (percentage) that contains inosine RNA modifications in total RNA (A) and ribosome-associated mRNA (B). C) Number of inosine RNA modified transcripts exhibiting one or multiple inosines per transcript in total RNA and ribosome-associated RNA. Numbers above line indicate the number of inosines / transcript. D) Number of inosine RNA modifications within specific regions (5’ UTR, CDS, intron, and 3’ UTR) in mRNA in total RNA and ribosome-associated RNA. *Means ± SEM within panel A and B are different (p < 0.05); significance was determined using 2 tests. Only transcripts with TPM ≥ 1 were analyzed.

### Consequences of coding sequence inosine RNA modifications in mouse oocytes

To understand the consequence of inosine RNA modifications, if any, on the protein coding capacity of mRNA transcripts, we used Ensembl Variant Effect Predictor (VEP) (McLaren et al. 2016) to identify the synonymous and non-synonymous substitutions (Fig. 3a). Among the non-synonymous substitutions, altered stop codons (stop loss, stop gain, or stop retained) made up less than 0.3% of coding sequence modifications. Therefore, stop codon substitutions were not included in further analyses.

**Figure 3.**
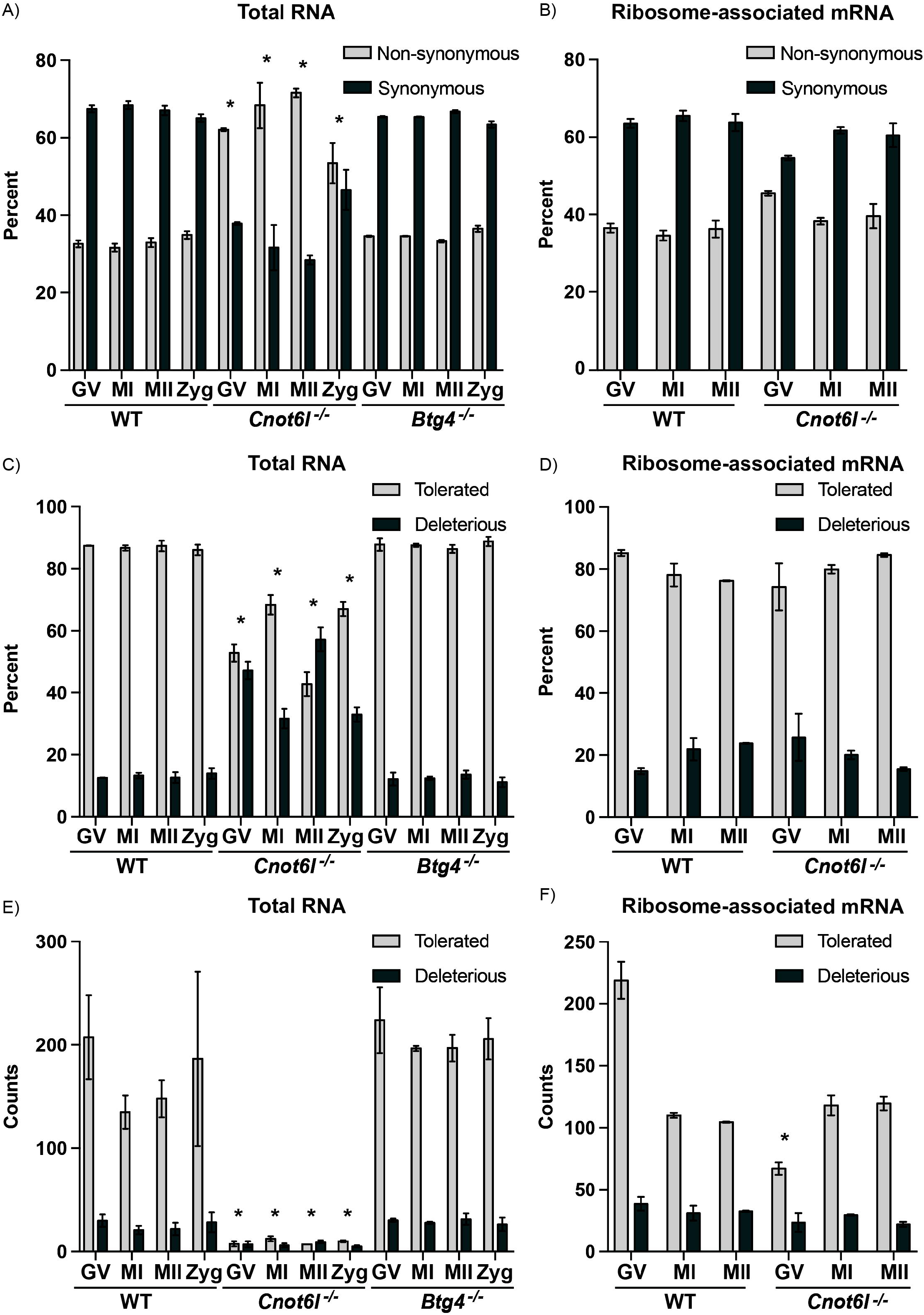
Consequences of inosine RNA modifications within CDS from WT, *Cnot6l^-/-^*, and *Btg4^-/-^* oocytes and zygote transcripts. A-B) Proportion of inosine modifications (percentage) in WT, *Cnot6l^-/-^*, and *Btg4^-/-^* oocytes and zygotes that result in non-synonymous or synonymous changes was determined for all CDS inosine modified mRNA transcripts in total RNA (A) and ribosome-associated mRNA (B). C-D) Proportion of tolerated and deleterious transcripts following Sorting Intolerant From Tolerant (SIFT) analysis of the inosine modified mRNA transcripts from WT, *Cnot6l^-/-^*, and *Btg4^-/-^* oocytes and zygotes in total RNA (C) or ribosome-associated mRNA (D). E-F) Counts of tolerated and deleterious transcripts resulting from inosine RNA modifications from WT, *Cnot6l^-/-^*, and *Btg4^-/-^* oocytes and zygotes in total RNA (E) or ribosome-associated mRNA (F). *Means ± SEM within a panel are different (p < 0.05); significance was determined using X2 tests or 2-way ANOVA. Only transcripts with TPM ≥ 1 were analyzed.

The majority of coding region inosine RNA modifications observed resulted in synonymous substitutions in WT and *Btg4^-/-^* oocytes (Brachova et al. 2019) Conversely, in oocytes from *Cnot6l^-/-^* oocytes, the pattern of inosine RNA modifications was reversed, with more than 60% exhibiting non-synonymous substitutions (Fig. 3a). Inosine RNA modified transcripts in the ribosome-associated mRNA had a similar pattern between WT and *Cnot6l^-/-^* oocytes (Fig. 3b). In order to predict the consequence of the amino acid substitutions, a computational tool, Sorting Intolerant From Tolerant (SIFT) that predicts the effects of amino acid substitution on protein function was used (Ng and Henikoff 2003). SIFT analysis showed that inosine RNA modifications observed in WT and *Btg4^-/-^* oocytes exhibited greater levels of tolerated amino acid substitutions when compared to the oocytes from *Cnot6l^-/-^* mice (Fig. 3c). Again, inosine RNA modified transcripts in the ribosome-associated mRNA had a similar pattern of tolerated amino acid substitutions between WT and *Cnot6l^-/-^* oocytes (Fig. 3d). When considering the number of tolerated or deleterious substitutions, *Cnot6l^-/-^* oocytes have reduced numbers in total RNA (Fig. 3e), but in the ribosome-associated transcripts, only GV *Cnot6l^-/-^* oocytes had significantly fewer tolerated substitutions (2-way ANOVA, p < 0.05, Fig. 3f). In summary, the total RNA fraction in *Cnot6l^-/-^* oocytes had a higher proportion of inosine RNA modifications within the coding sequencing that resulted in potentially deleterious non-synonymous substitutions. However, within the ribosome-associated RNA, similar levels of synonymous and non-synonymous and tolerated and deleterious substitutions were observed between WT and *Cnot6l^-/-^*. These results suggest that translated mRNA in WT and *Cnot6l^-/-^* oocytes share a similar capacity for inosine induced protein recoding through alterations to codon usage.

### Inosine RNA modifications are enriched at the wobble position in ribosome-associated RNA

The abundance of inosine modifications resulting in synonymous substitutions in WT and *Btg4^-/-^* oocyte RNA led us to investigate the potential effects of inosine RNA modifications on codon usage. We first determined the number of inosine RNA modifications that occur in the 34 different codons that contain an adenosine, excluding stop codons. We found in WT and *Btg4^-/-^* oocytes, certain codons contain more inosine RNA modifications: AAA, ACA, CAA, CCA, GAA, and GCA (Fig. 4a). In *Cnot6l^-/-^* oocytes, inosine-modified codons were significantly reduced compared with WT and *Btg4^-/-^* oocytes (Fig. 4a). Additionally, there were no enriched modified codons detected in *Cnot6l^-/-^* oocytes (Fig. 4a). Within transcripts from ribosome-associated mRNA in both WT and *Cnot6l^-/-^* oocytes, we detected similar codons enriched in inosine RNA modifications (Fig. 4b). Further analysis indicated that inosine modifications are enriched at the wobble position in codons with multiple adenosines in total RNA of WT (Fig. 4c), but not in *Cnot6l^-/-^* oocytes. Conversely, both WT and *Cnot6l^-/-^* ribosome-associated mRNA exhibited the wobble position enhancement (Fig. 4d). The wobble position RNA modification across all codons was not present in *Cnot6l^-/-^* oocytes in total RNA, in contrast to WT and *Btg4^-/-^* oocytes, while the ribosome-associated mRNA retained wobble position inosine RNA modifications across all three genotypes (Fig. 4e and f). Other codons assessed did not show a specific enrichment or difference between WT and *Cnot6l^-/-^* oocytes (Supplemental Fig. S2).

**Figure 4.**
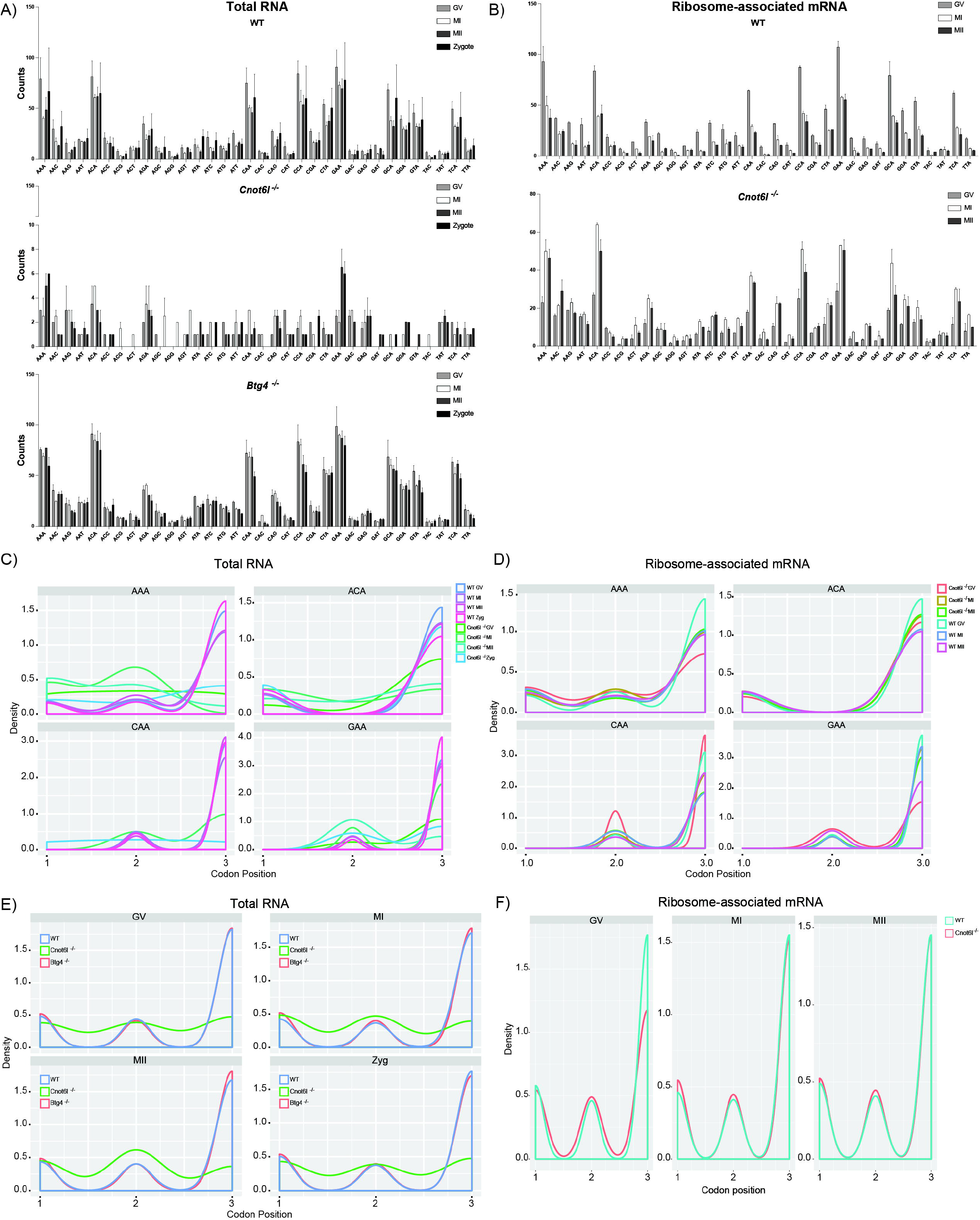
Inosines are enriched at the wobble position of codons in total RNA and ribosome-associated RNA. A-B) Number of inosine RNA modifications in adenosine-containing codons in WT, *Cnot6l^-/-^*, and *Btg4^-/-^* oocytes and zygotes in total RNA (A) and ribosome-associated RNA (B). C-D) Number of inosine RNA modifications occurring at specific codons among the codons with multiple adenosines in WT, *Cnot6l^-/-^*, and *Btg4^-/-^* oocytes and zygotes in total RNA (C) and ribosome-associated RNA (D). E-F) Global frequency of inosine RNA modifications at the first, second, or third codon positions in WT, *Cnot6l^-/-^*, and *Btg4^-/-^* oocytes and zygotes in total RNA (E) and ribosome-associated RNA (F). *Means ± SEM within a panel were different (p < 0.05); significance was determined using two-way ANOVA tests. Only codons from transcripts with TPM ≥ 1 were analyzed.

### Efficiency of inosine RNA modifications is highest in ribosome-associated mRNA

Transcriptome-wide inosine RNA modification efficiency at various codons remained around 50% throughout oocyte maturation and in the zygote in WT oocytes (Fig. 5a, raw counts are in Supplemental Fig. 3). Conversely, *Cnot6l^-/-^* oocytes displayed defects in inosine RNA modification efficiency in total RNA at all stages of oocyte maturation and in zygotes (Fig. 5a). An examination of inosine RNA modification efficiency in *Btg4^-/-^* oocytes revealed a reduction of modification efficiency of select codons (Fig. 5a). The inosine RNA modification efficiency reached almost 100% within transcripts of ribosome-associated mRNA of WT oocytes, and then decreased during oocyte maturation (Fig. 5b). *Cnot6l*^-/-^ oocytes also displayed almost 100% efficiency of particular codons (AAA, AAC, AAT, AGA, CAA, CAG, CCA, CGA, GTA, TTA), however other codons were not modified at all (AAG, ACG, ACT, AGG, ATT, GAC, GAG, GAT), as seen in Fig. 4. By the end of oocyte maturation, the inosine RNA modification efficiency was completely lost (Fig. 5b). Overall, during oocyte maturation, the inosine RNA modification efficiency is almost 100% in ribosome-associated mRNA, and decreases during maturation. These results suggest that, in the *Cnot6l^-/-^* oocytes where mRNA is stabilized, the inosine modified mRNA are degraded, while the unmodified mRNA stabilized.

**Figure 5.**
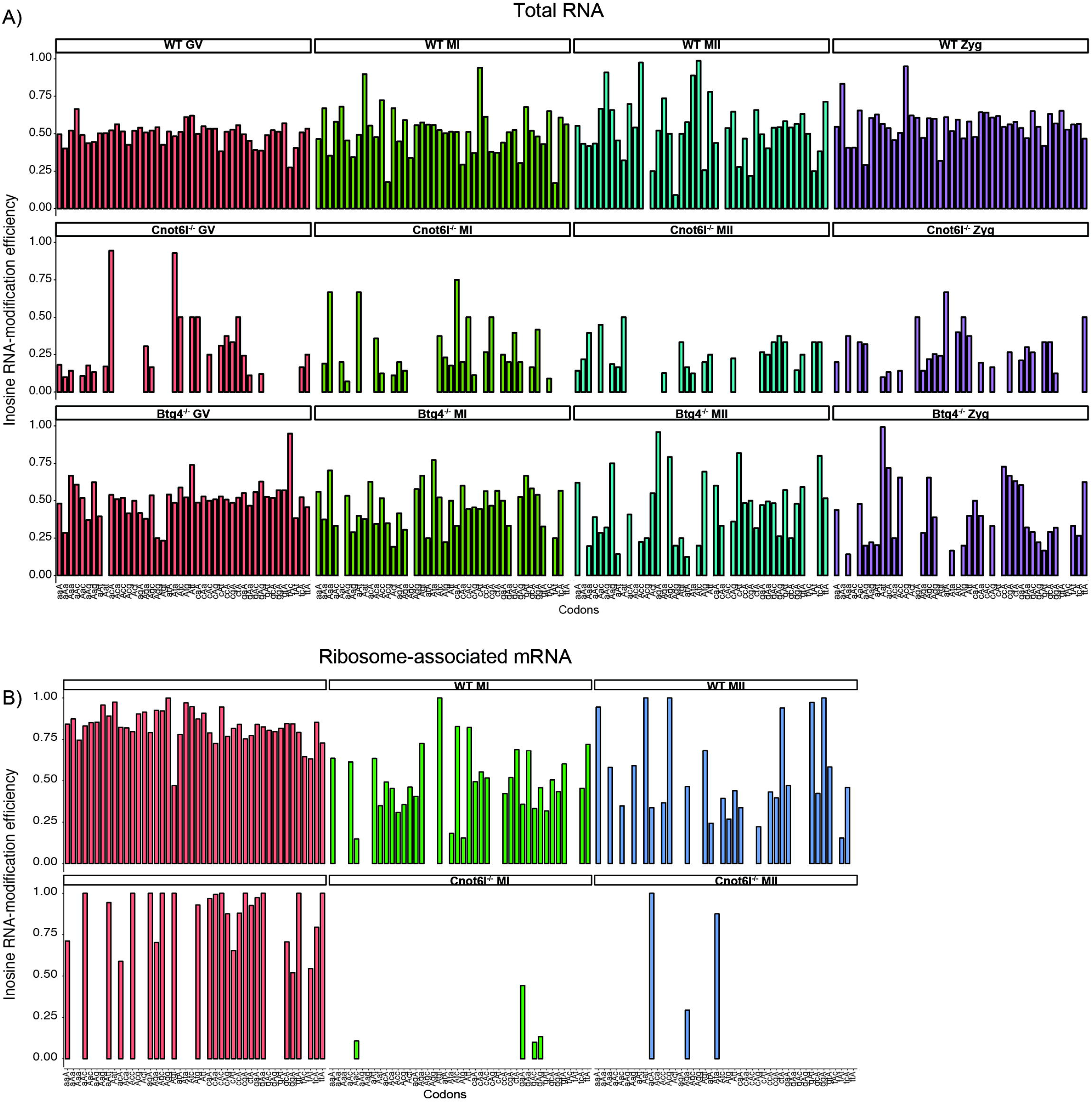
Inosine RNA modification efficiency of total and ribosome-associated mRNA. A) Inosine RNA modification efficiency in total RNA from WT, *Cnot6l^-/-^*, and *Btg4^-/-^* oocytes and zygotes. Capital letter denotes the position of the codon with an inosine RNA modification. B) Inosine RNA modification efficiency in ribosome-associated mRNA from WT and *Cnot6l^-/-^* oocytes and zygotes. Capital letter denotes the position of the codon with an inosine RNA modification.

## Discussion

Translational mRNA decay is a main mechanism of maternal mRNA clearance in mammalian oocytes. During oocyte maturation, mRNA are recruited for translation and subsequent degradation through interactions with components of the CCR4-NOT complex (Sha et al. 2018; Vieux and Clarke 2018). The maternal-to-zygotic decay machinery component CNOT6L plays an important role in maternal mRNA decay during MI (Sha et al. 2018; Vieux and Clarke 2018; Horvat et al. 2018), while the adaptor factor BTG4 plays an important role in degradation of maternal mRNA after ovulation and prior to fertilization (Liu et al. 2016; Yu et al. 2016). As such, selective decay of transcripts allows for specific control of events occurring during the MZT, necessary for early embryo development.

An important component of translational mRNA decay is the rate of translation, which is influenced by codon composition, a phenomenon known as codon optimality (Presnyak et al. 2015; Bazzini et al. 2016; Carneiro et al. 2019; Horstick et al. 2015; Pop et al. 2014; Bergman and Tuller 2020). Codons that are non-optimal slow the ribosome, and experimental evidence indicates that this increases mRNA degradation through CCR4-NOT (Webster et al. 2018). It has recently been reported that inosine RNA modifications in the coding sequence of mRNA lead to ribosome stalling (Licht et al. 2019). In a previous report (Brachova et al. 2019), and now here in WT and *Btg4^-/-^* oocytes, we identified inosine RNA modifications specifically enriched in at the codon wobble position in oocytes (Brachova et al. 2019). Based on our finding and others, we hypothesized that inosine RNA modifications within the coding sequence could alter mRNA stability, potentially through translational mRNA decay. To test this hypothesis we analyzed previously generated RNA-seq data for *Cnot6l^-/-^* and *Btg4^-/-^* oocytes, which are known to exhibit dysfunction in translational mRNA decay (Sha et al. 2018). The nearly complete loss of inosine RNA modified transcripts in the *Cnot6l^-/-^* oocytes was unexpected, as we originally hypothesized to identify increased modified transcripts in oocytes with hyper-stabilized mRNA as previously established (Sha et al. 2018). The loss of inosine RNA modifications was not associated with a loss of *Adar* expression (Supplemental Fig. S4), but we have not evaluated ADAR protein levels in *Cnot6l^-/-^* oocytes, which is an important consideration. *Btg4^-/-^* oocytes, on the other hand, have inosine RNA modification profiles similar to that of WT oocytes. Both CNOT6L and BTG4 accumulate during oocyte maturation and trigger maternal mRNA decay, but their knockouts result in different oocyte phenotypes: *Cnot6l^-/-^* oocytes arrest at prometaphase I, while *Btg4^-/-^* oocytes are able to complete meiotic maturation and do not arrest until fertilization (Sha et al. 2018). CNOT6L and BTG4 degradation pathways act on similar substrate pools during oocyte maturation, albeit with different timing (Sha et al. 2018). We identified inosine modified transcripts common between all genotypes, and observed that *Cnot6l^-/-^* oocytes had significantly fewer modified transcripts compared to WT and *Btg4^-/-^* oocytes (Fig. 1c). We consider three possible mechanisms that can explain our results. First, CONT6L and ADAR could interact in oocytes, facilitating inosine RNA modifications. Biological evidence for such a complex is lacking. Recent mass spectrometry approaches to identify ADAR associated proteins that regulate inosine RNA modifications have not identified CCR4-NOT complex components (Freund et al. 2020). Rather, several components of the ribosome were identified directly associated with ADAR (Freund et al. 2020).

The second explanation for reduced inosine RNA modifications in *Cnot6l^-/-^* oocytes is that *Adar* is no longer being translated efficiently and the ADAR protein level is reduced, thus reducing inosine RNA modifications. Though RNA abundance is not direct evidence for translation, we detected *Adar* transcripts in *Cnot6l^-/-^* oocytes in both the total RNA and ribosome-associated fractions (Supplemental Fig. S4). Additionally, it has been reported that *Cnot6l^-/-^* oocytes over-translate mRNA (Sha et al. 2018). Based on these observations, we would anticipate that *Adar* would have a higher probability to be over-translated rather than under-translated. Furthermore, analysis of the ribosome-associated mRNA revealed that the number of inosine modified transcripts were similar between MI and MII stages in WT and *Cnot6l^-/-^* oocytes (Fig. 1b, d). This indicates that within the ribosome-associated fraction, ADAR had similar levels of activity in WT and *Cnot6l^-/-^* oocytes. We observed a significant difference in inosine modified mRNA between WT and *Cnot6l^-/-^* GV oocytes (Fig. 1b, d), indicating the reduction of inosine RNA modifications occurs prior to GV entry. Additionally, analysis of the number of inosine modifications per transcript, codon wobble position modifications, and efficiency of modifications within ribosome-associated mRNA revealed that both WT and *Cnot6l^-/-^* oocytes had similar levels of inosine RNA modifications at the GV stage (Fig. 3c, 4f, & 5b). We have previously shown that the inosine modification efficiency in oocytes is linked to the catalytic activity of ADAR (Brachova et al. 2019), indicating that ADAR is present and active in *Cnot6l^-/-^* oocytes. Therefore, we do not consider a reduction of ADAR protein to be a likely explanation for the decrease in inosine RNA modifications present in *Cnot6l^-/-^* oocytes, however, further experimentation would be needed to test ADAR levels in the absence of CNOT6L.

A third possible mechanism to explain the reduction in inosine RNA modifications in *Cnot6l^-/-^* oocytes is that CONT6L and ADAR both contribute to translational mRNA degradation, through two independent, but parallel pathways. An important component of translational mRNA decay is the rate of translation, which is influenced by codon composition, a phenomenon known as codon optimality (Presnyak et al. 2015; Bazzini et al. 2016; Carneiro et al. 2019). Codons that are non-optimal slow the ribosome, and experimental evidence indicates that this increases mRNA degradation through CCR4-NOT (Webster et al. 2018; Buschauer et al. 2020). It has been reported that inosine RNA modifications in the coding sequence of mRNA lead to ribosome stalling (Licht et al. 2019). In our previous report, we identified inosine RNA modifications in the total RNA fraction that are specifically enriched at the codon wobble position (Brachova et al. 2019). Here, we show that ribosome-associated mRNA have similar inosine enrichment to the wobble position of codons in both WT and *Cnot6l^-/-^* oocytes (Fig. 4f). However, codon wobble position enrichment of inosine is absent in the total RNA fraction *Cnot6l^-/-^* oocytes (Fig. 4e). Furthermore, the number of transcripts with inosine RNA modifications is significantly reduced in the total RNA fraction of *Cnot6l^-/-^* oocytes when compared to WT (Fig. 1a). We reason that one explanation for the difference between the amount of RNA transcripts with inosine RNA modifications between total RNA and ribosome-associated mRNA fractions is due to the number of oocytes used and the subsequent enrichment of ribosome fraction (total RNA: 10 oocytes; ribosome-associated RNA: 500 oocytes). The inosine modified transcripts in the ribosome fraction are also present in the total RNA, albeit at a smaller proportion, and enrichment allows for their detection.

Additionally, the observed reduction of inosine RNA modifications in *Cnot6l^-/-^* oocytes within the total RNA fraction may be due to two factors: overall increased mRNA stability and selective decay of inosine modified mRNA. In *Cnot6l^-/-^* oocytes, mRNA is globally stabilized, and we hypothesize that inosine modified RNA is targeted for decay. We reason that the stabilization of mRNA increases the mRNA complexity of the total RNA fraction, thus reducing the sensitivity of RNA-seq to detect inosine modified RNAs. The same phenomenon is present in the ribosome-associated fraction of *Cnot6l^-/-^* oocytes, but it is offset by the enrichment procedure of ribosome. In the ribosome-associated fraction, we were able to observe that in the GV oocyte stage of WT and *Cnot6l^-/-^* oocytes, inosine RNA modification efficiency was above 60%, and in some cases almost 100% for the codons analyzed (Fig. 5b). In comparison, the total RNA fraction, inosine modification efficiency was approximately 50% (Fig. 5a). In the ribosome-associated fraction of WT oocytes, progression into MI phase was marked by a reduction in inosine RNA modification efficiency (Fig. 5b). In *Cnot6l^-/-^* oocytes, there is almost a complete loss of inosine RNA modification efficiency at the codons analyzed (Fig. 5b). An interpretation of this observation is that inosine mRNA are subject to a decay process independent of CNOT6L. In ribosome-associated mRNA of WT oocytes, we observe a reduction of inosine containing mRNA, and this reduction is similar to the reduction in overall mRNA levels at MI (Sha et al. 2018). We hypothesize that mRNA decay mediated by CNOT6L and inosine RNA modifications share a common component: translation. It has previously been shown that inosine RNA modifications in coding sequences can induce ribosome stalling (Licht et al. 2019). Stalling, or slowing of ribosomes can induce CCR4-NOT decay (Webster et al. 2018; Buschauer et al. 2020). Based on this prediction, we had originally hypothesized that we would observe increases in inosine RNA modifications in *Cnot6l^-/-^* oocytes. However, we observed a decrease in inosine RNA modifications in the ribosomal associated fraction. If the inosine induced stalling or slowing of ribosomes is still occurring and in the absence of a functioning CCR4-NOT complex decay was still progressing, this would indicate a parallel pathway to mRNA decay. Our results suggest a novel mechanism of mRNA clearance during oocyte maturation, in which inosine-containing transcripts decay uncoupled from CCR4-NOT deadenylation. Further experiments will reveal the impact of inosine RNA modifications on maternal mRNA decay.

## Methods

### Sources of GV and MII RNA-seq datasets

Wild-type GV, MI, MII, and Zyote RNA-seq data, and GV, MI, MII, and Zygote RNA-seq data from *Cnot6l^-/-^* and *Btg4^-/-^* knockout mice dataset PRJNA486094 (Sha et al. 2018). The authors of this study generated library preparations from total RNA isolated from groups of 10 oocytes or embryos, all on the C57B6 background. For RNA-seq libraries of polysome-bound mRNA, groups of 500 oocytes or embryos were used for polysome pull-down. Two replicates were generated for library preparation, and each preparation had mCherry spike-in added before cDNA synthesis for normalization. Barcoded libraries were pooled and sequenced on an Illumina HiSeq X Ten platform with 150 bp paired-end reads (Sha et al. 2018).

### Identification and consequence analysis of inosine RNA modifications

We identified putative inosine RNA modifications utilizing a combination of the HISAT2 aligner, the Genome Analysis Tool Kit (GATK), and Ensembl Variant Effect Predictor (VEP) (McKenna et al. 2010; McLaren et al. 2016; Kim et al. 2015; Brachova et al. 2019). Inosines appear as A-to-G substitutions when comparing RNA-seq data to a reference genome (Ramaswami et al. 2013). HISAT2 was used to map raw RNA-seq reads to a mus musculus index built from GRCm38 containing common SNP annotation from dbSNP.

Default settings for HISAT2 were used for each type of library aligned (i.e. single-end, paired-end, or stranded, Supplemental Table 2) (Kim et al. 2015). Prior to alignment, fastq files were checked for adaptors and trimmed if necessary, using Trimmomatic (Bolger et al. 2014). The default alignment settings for HISAT2 will report at most 10 valid primary alignments; we did not increase the amount of multimapping allowed. HISAT2 reports valid alignments that are at or above the minimum alignment score determined by the function where the length of the read. Mismatches in the alignment were subtracted from the alignment score, reads that fell below the minimum alignment score were not reported. RNA/DNA differences were called using the Genome Analysis Toolkit (GATK) RNASeq variant pipeline with modifications (McKenna et al. 2010). The program elprep was used to sort, mark duplicates, and index RNA-seq reads. Known SNPs were filtered out using the Mouse Genomes Project database, Wellcome Trust Sanger Institute mouse strains (Yalcin et al. 2011). RNA-DNA mismatches identified by HISAT2, regardless if stranded mode was used, and GATK were reported in the sense orientation; therefore strand information of the variant is inferred from the gene model using Ensembl VEP (McLaren et al. 2016). VCF files were filtered on allelic depth (AD > 0) and used as input for Ensembl Variant Effect Predictor (VEP, (McLaren et al. 2016)). VEP was used to identify inosine modified transcripts, categorize the location within the transcript; and determine the consequence of inosines on coding capacity. Only inosine sites occurring in transcripts with TPM ≥ 1 and having an AD > 0 were reported. A transcript was considered to be inosine modified if it contained at least one inosine site. R statistical computation software with the following packages was used to parse VEP output: sleuth, biomaRt, dplyr, plyr, AnnotationFuncs, org.Mm.eg.db, ggplot2 (Pimentel et al. 2017; Team and Others 2013; Wickham 2009; Durinck et al. 2009).

## Acknowledgements

This work was supported by the National Institutes of Health (HD094545 to L.K.C., HD099269, CA200357 to P.B., and HD094545-01A1S1 to N.S.A.). The authors also acknowledge the University of Kansas Medical Center Internal Support and Genomics Core and their funding: Kansas Intellectual and Developmental Disabilities Research Center (NIH U54 HD090216).

