## Supplemental Figure S1 for "Loss of *Cnot6l* impairs inosine RNA modifications in mouse oocytes"

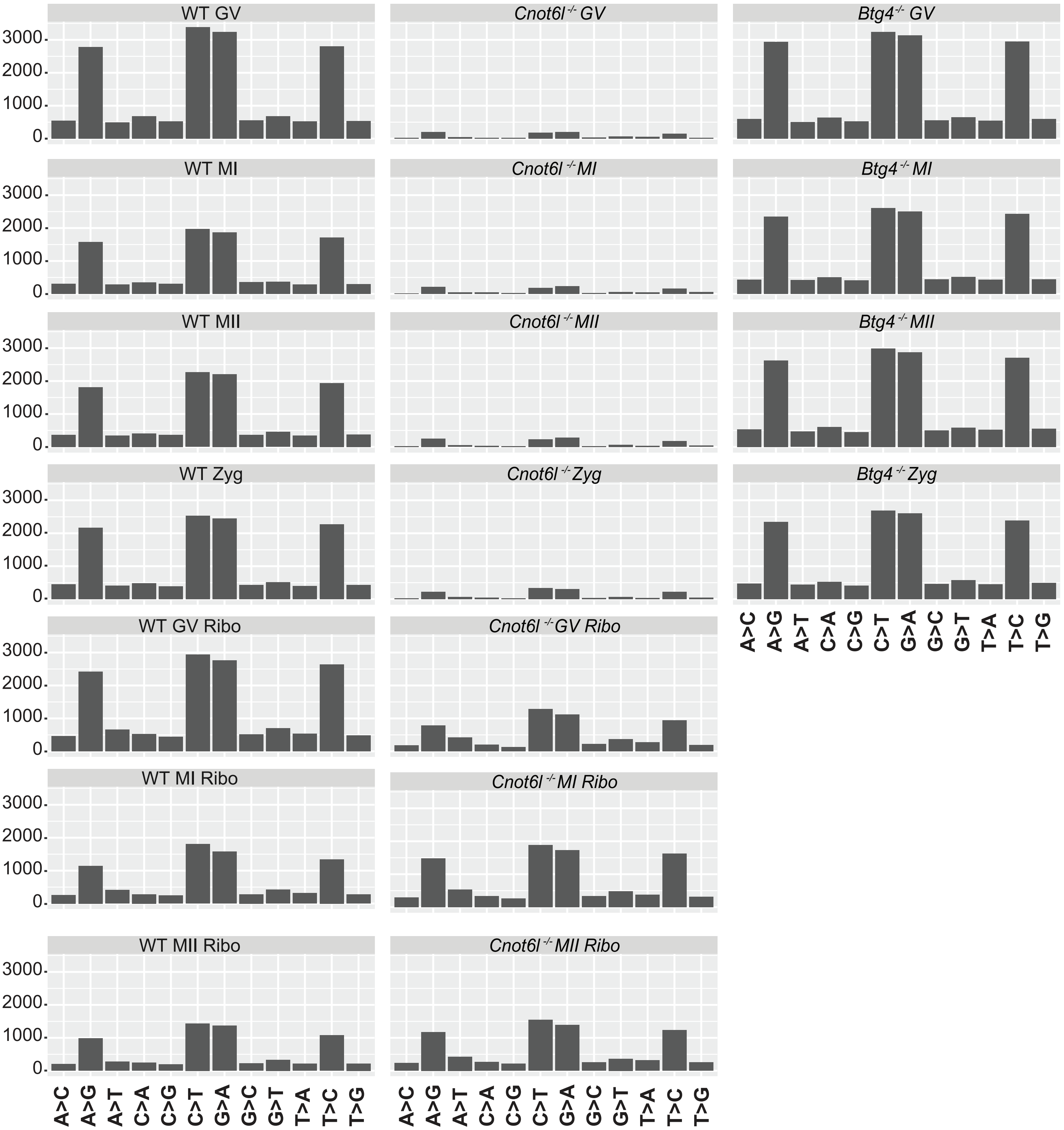


Supplemental Figure S1. Identified single nucleotide substitutions in GV or MI oocytes, or MII eggs from WT, Cnot6l$^{-/-}$ total RNA and ribosome-associated RNA, and Btg4$^{-/-}$ total RNA. Single nucleotide substitution counts identified by our RNA modification discovery pipeline in the 5′ UTR, CDS, intron, and 3′ UTR.
