## Supplemental Figure S2 for "Loss of *Cnot6l* impairs inosine RNA modifications in mouse oocytes"

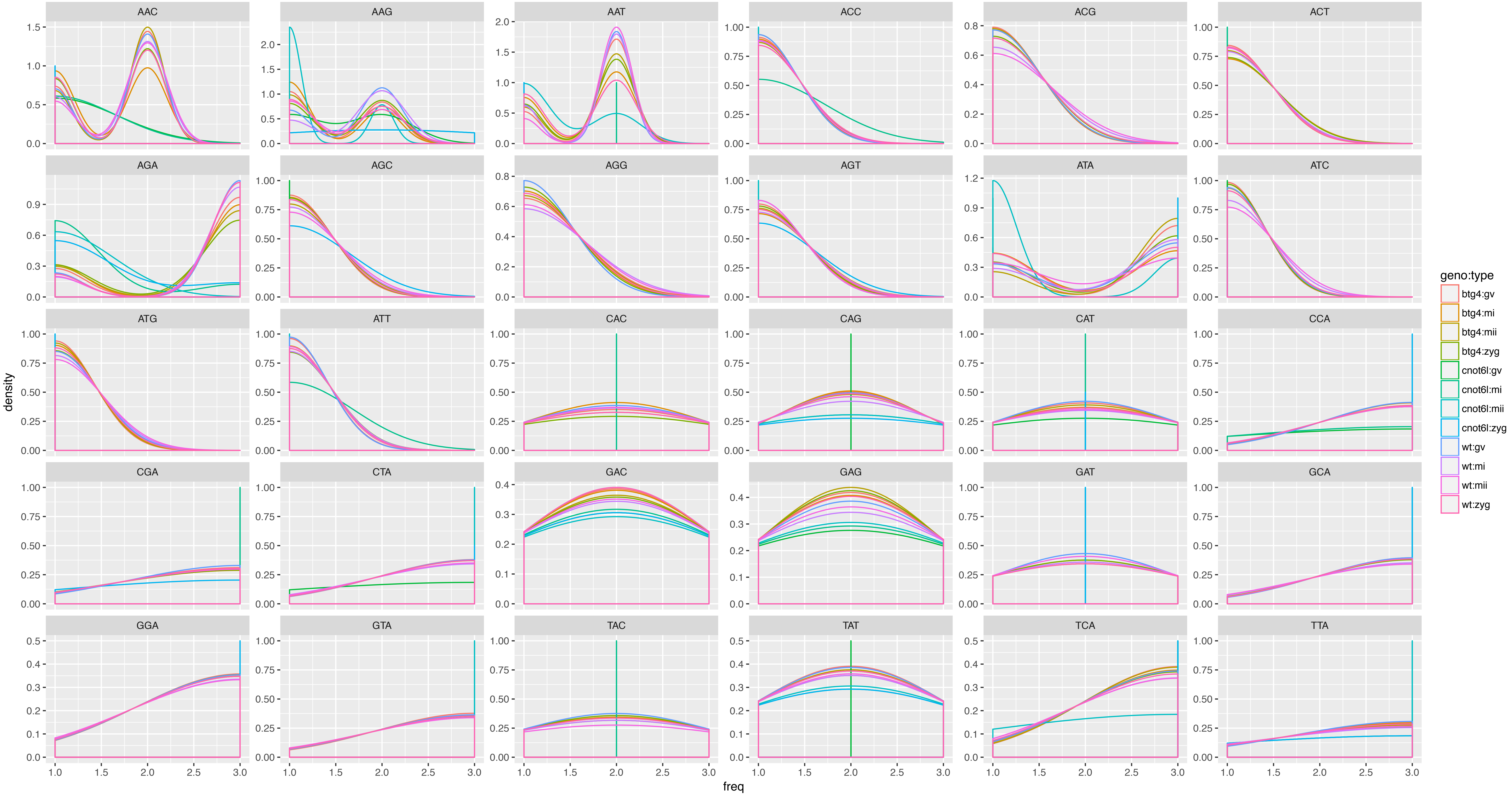


Supplemental Figure S2. Codon usage in total RNA from WT, Cnot6l-/-, and Btg4-/- oocytes and zygotes. The frequency of codons occurring in all transcripts with a TPM ≥ 1.
