## Supplemental Figure S3 for "Loss of *Cnot6l* impairs inosine RNA modifications in mouse oocytes"

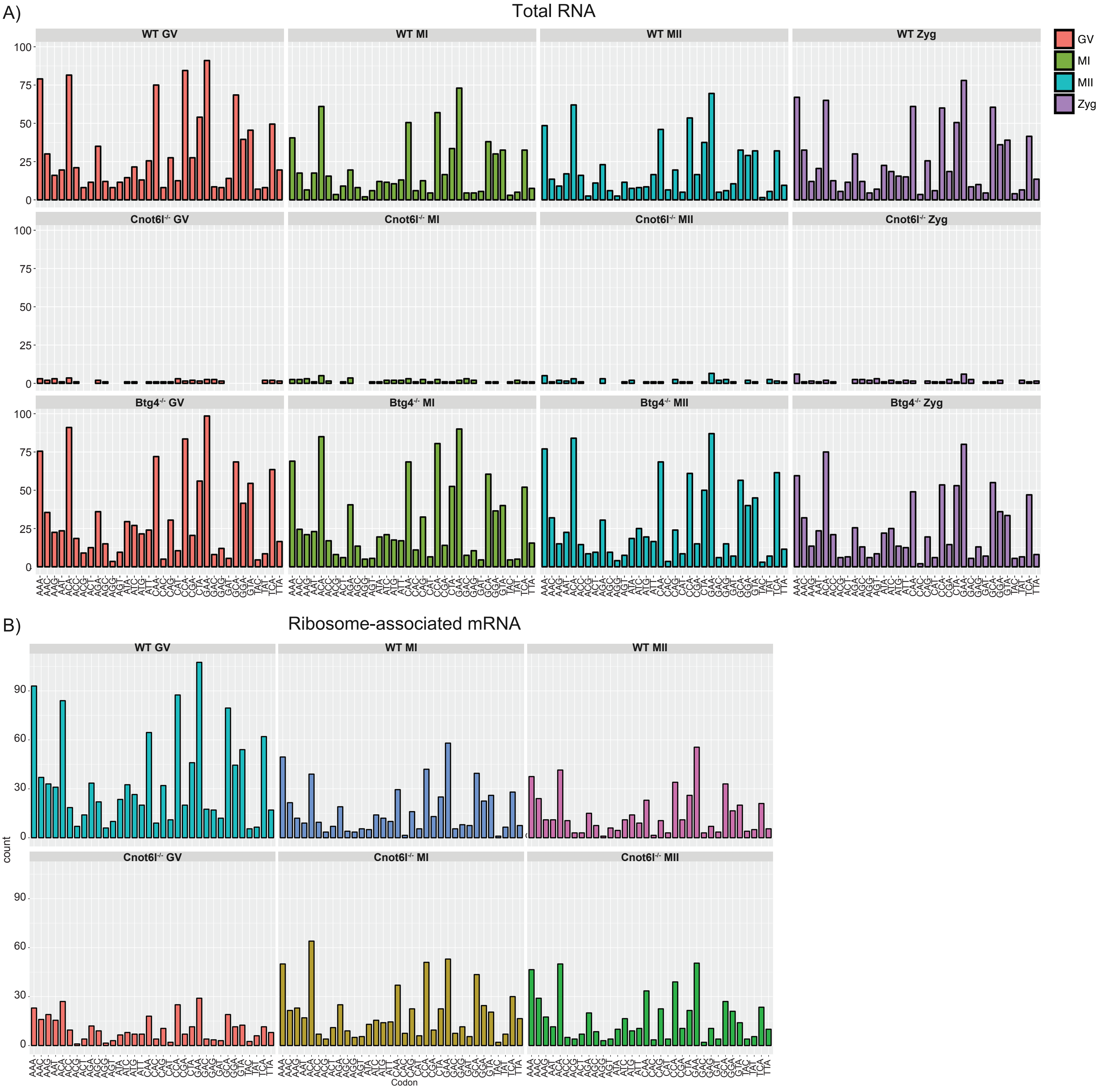


Supplemental Figure S3. Counts of inosine RNA modifications in total RNA and ribosome-associated mRNA. A) Inosine RNA modifications in total RNA from WT, Cnot6l$^{-/-}$, and Btg4$^{-/-}$ oocytes and zygotes. B) Counts of inosine RNA modifications in ribosome-associated mRNA from WT and Cnot6l-/- oocytes and zygotes.
