## Supplemental Figure S4 for "Loss of *Cnot6l* impairs inosine RNA modifications in mouse oocytes"

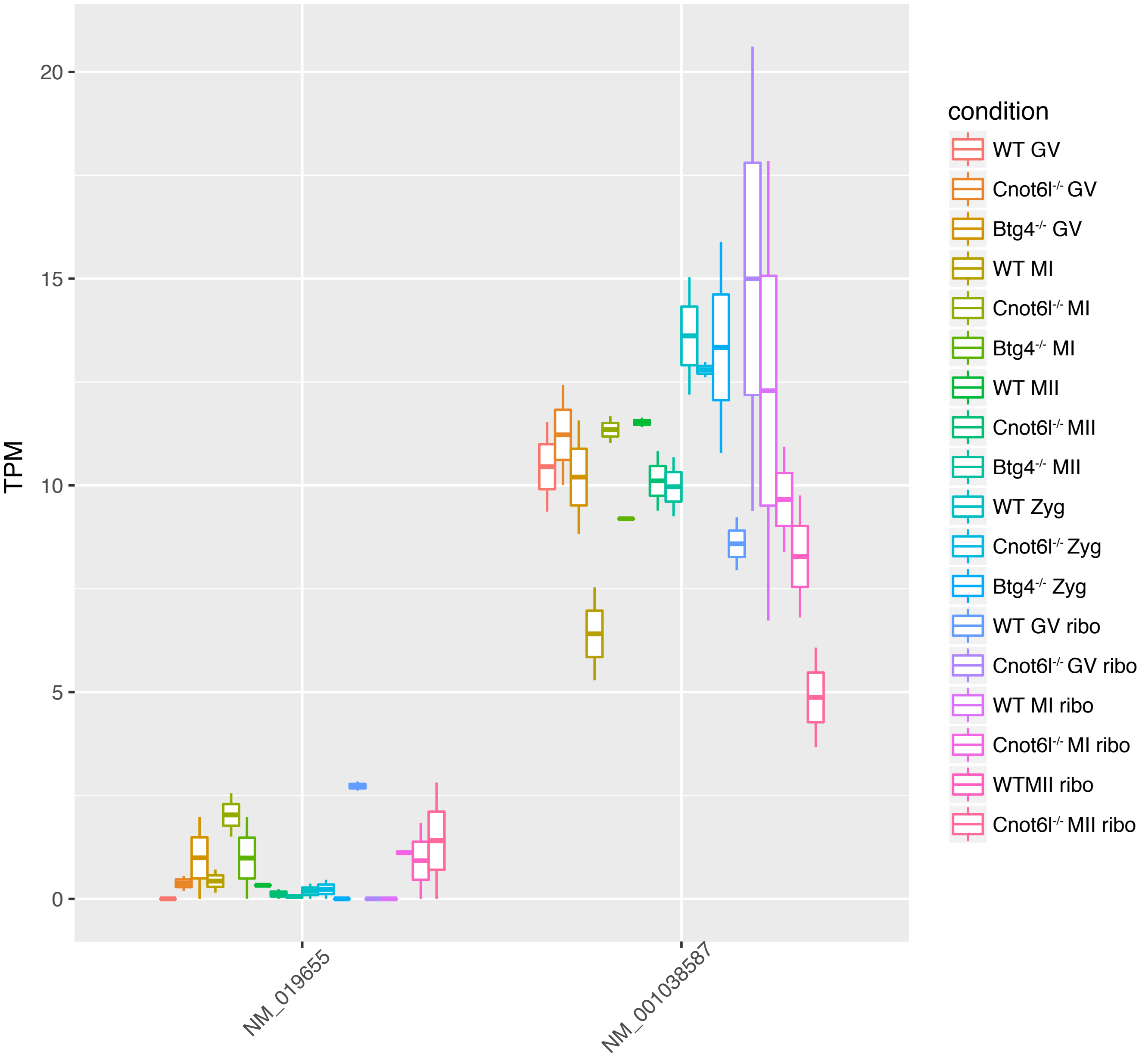


Supplemental Figure S4. Adar isoform abundance in total RNA and ribosome-associated mRNA from WT, Cnot6l-/-, and Btg4-/- oocytes and zygotes. TPM= transcripts per million.
